# mRNA-based vaccine candidate COReNAPCIN^®^ induces robust humoral and cellular immunity in mice and non-human primates

**DOI:** 10.1101/2022.03.05.483092

**Authors:** Reza Alimohammadi, Meysam Porgoo, Mohamad Eftekhary, Seyed Hossein Kiaie, Ehsan Ansari Dezfouli, Maryam Dehghani, Kaveh Nasrollahi, Talieh Malekshahabi, Maryam Heidari, Sedigheh Pouya, Masoumeh Alimohammadi, Dorsa Sattari Khavas, Mohammad Sadra Modaresi, Mohammad Hossein Ghasemi, Hamed Ramyar, Fatemeh Mohammadipour, Fateme Hamzelouei, Ahmadreza Mofayezi, Seyed Saeed Mottaghi, Amirhosein Rahmati, Mohsen Razzaznian, Vista Tirandazi, Fatemeh Borzouee, Hossein Sadeghi, Melika Haji Mohammadi, Leila Rastegar, Seyed Milad Safar Sajadi, Hossein Ehsanbakhsh, Hamed Bazmbar, Maedeh Shams Nouraee, Pouya Pazooki, Mina PahlevanNeshan, Khadijeh Alishah, Fateme Nasiri, Neda Mokhberian, Seyedeh Shima Mohammadi, Shima Akar, Hamidreza Niknam, Marzyieh Azizi, Mohammad Ajoudanian, Mohammad Hossein Moteallehi-Ardakani, Seyed Ali Mousavi Shaegh, Reihaneh Ramezani, Vahid Salimi, Reza Moazzami, Seyed Mahmoud Hashemi, Somaye Dehghanizadeh, Vahid Khoddami

## Abstract

At the forefront of biopharmaceutical industry, the messenger RNA (mRNA) technology offers a flexible and scalable platform to address the urgent need for world-wide immunization in pandemic situations. This strategic powerful platform has recently been used to immunize millions of people proving both of safety and highest level of clinical efficacy against infection with severe acute respiratory syndrome coronavirus 2 (SARS-CoV-2). Here we provide preclinical report of COReNAPCIN^®^; a vaccine candidate against SARS-CoV-2 infection. COReNAPCIN^®^ is a nucleoside modified mRNA-based vaccine formulated in lipid nanoparticles (LNPs) for encoding the full-length prefusion stabilized SARS-CoV-2 spike glycoprotein on the cell surface. Vaccination of C57BL/6 and BALB/c mice and rhesus macaque with COReNAPCIN^®^ induced strong humoral responses with high titers of virus-binding and neutralizing antibodies. Upon vaccination, a robust SARS-CoV-2 specific cellular immunity was also observed in both mice and non-human primate models. Additionally, vaccination protected rhesus macaques from symptomatic SARS-CoV-2 infection and pathological damage to the lung upon challenging the animals with high viral loads of up to 2×10^8^ live viral particles. Overall, our data provide supporting evidence for COReNAPCIN^®^ as a potent vaccine candidate against SARS-CoV-2 infection for clinical studies.

## 1. Introduction

Severe acute respiratory syndrome coronavirus 2 (SARS-CoV-2) is a member of the beta coronavirus genus, and since its emergence in 2019 leads to hundreds of millions of cases of Coronavirus Disease 2019 (COVID-19) and several millions of deaths so far^1,2^. Unlike other members of beta coronavirus genus which led to substantial human disease in the last 20 years, SARS-CoV-2 has airborne transmission effectively spreading between individuals^3,4^. Despite of rapid development of vaccines against SARS-CoV-2, there is still a substantial worldwide shortage of effective vaccines, especially with rapid emergence and global spread of new SARS-CoV-2 variants^5^.

In the past two years several vaccine platforms have been developed to prevent SARS-CoV-2, among which the mRNA technology, due to some unique features, appears to be the most efficient platform^6^. Notably the high-speed, scalable and flexible production of mRNA vaccines is critical for the rapid response to pandemic conditions. Only few weeks can take from receiving the genetic code of a particular antigen to the design and production of mRNA vaccines in a prepared platform, an important feature especially for generation of variant-specific vaccines in the case of COVID-19^7–10^. Beside unique advantages in the development and mass production phases, the mRNA vaccines have been shown to generate the most efficient and safest vaccines^11^. This is primarily due to the in-vivo translation of mRNAs into the target protein, ensuring the correct folding and natural glycosylation pattern of the encoded antigen. Additionally, both humoral and cellular immunity are efficiently elicited using mRNA vaccines, generating the most effective vaccines against viruses including SARS-CoV-2^12^. Notably, while formulation of mRNA vaccines is free of adjuvants, increasing their safety indices, the chemical nature of foreigner mRNA molecules entering the cells from outside confers a significant intrinsic adjuvant-like activity to mRNA vaccines efficiently enhancing both innate and adaptive immune responses. Moreover, due to their transient mode of action there is no concern about integration of mRNA codes into the genomic DNA^13^. Overall, the mRNA technology, recently proven by at least two approved and globally tested mRNA vaccines, introduces an efficient strategy to obtain the highest level of both humoral and cellular immunity against a particular infectious agent, in parallel with the highest safety indices, making it one of the best platforms for vaccine development.

SARS-CoV-2 Spike protein (S protein), the primary target for neutralizing antibodies, is a heavily glycosylated protein that orchestrates the entrance of the virus into the host cells. During cell fusion, the S protein undergoes conformational change and facilitates the insertion of virus genome into the host cells^14,15^. This change in the original 3 dimensional (3D) structure of S protein may have a catastrophic effect on S protein immunogenicity^16^. One way to increase the immunogenicity of S protein is the substitution of two proline residues at the apex of the central helix and heptad repeat 1 of the spike glycoprotein to stabilize it at its native prefusion conformation^17,18^.

Here we present the preclinical report of COReNAPCIN^®^, a modified mRNA vaccine candidate formulated in lipid nanoparticles (LNPs) for expression of the prefusion-stabilized form of SARS-CoV-2 spike glycoprotein upon intramuscular injection, in rodents and non-human primates.

## 2. Results

### 2.1. Design, production and characterization of SARS-CoV-2 Spike mRNA-LNP

To make COReNAPCIN^®^ the nucleoside modified mRNA encoding prefusion stabilized form of SARS-CoV-2 Spike glycoprotein was designed and produced. The 2P mutation was used to stabilize the spike protein in the prefusion conformation to increase the efficiency of immunologic response against viral infection (fig. 1a&b). The COReNAPCIN^®^ coding sequence was designed based on a special codon optimized template for maximizing the protein expression (by substituting the rare codons with common triplets) and minimizing the intracellular immunity (by reducing the uridine stretches within the sequence). Additionally for further stabilization of the mRNA transcripts and elimination of the unfavorable immunogenic responses all remaining uridines within the sequence were substituted with N1-methyl-pseudouridine during the mRNA synthesis step. After assessment of RNA integrity and purity (fig. 1c&g), the nucleoside modified mRNA was formulated into lipid nanoparticles (LNPs) by mixing the purified mRNAs with special lipids in a microfluidic device (fig. 1d) (see materials and methods).Evaluation of the resulting mRNA-LNPs by dynamic light scattering (DLS) showed average particle size of 104.3 nm, zeta potential (ZP) of -5.26, and polydispersity index (PDI) of 0.196 (fig. 1e), and transmission electron microscopy (TEM) further confirmed the uniformity in shape and size of nanoparticles and validated their expected LNP-specific morphology. RNA encapsulation efficiency (EE%) of mRNA-LNPs was analyzed semi quantitatively using gel retardation assay and quantitatively using a fluorescent-based method showing EE% of more than 90 (Table s1). For in-vitro functional analysis following addition of mRNA-LNPs to cultured HEK-293 cells the expression of SARS-CoV-2 Spike glycoprotein-2P at the cell surface was evaluated by flow cytometry using spike-specific antibodies (fig. 1h). Further validation of correct protein translation was performed by Western Blotting (fig. 1i). After full characterization COReNAPCIN^®^ was used for assessment of immunity in C57BL/6 and BALB/c mice and rhesus macaque (see below).

**Figure 1:**
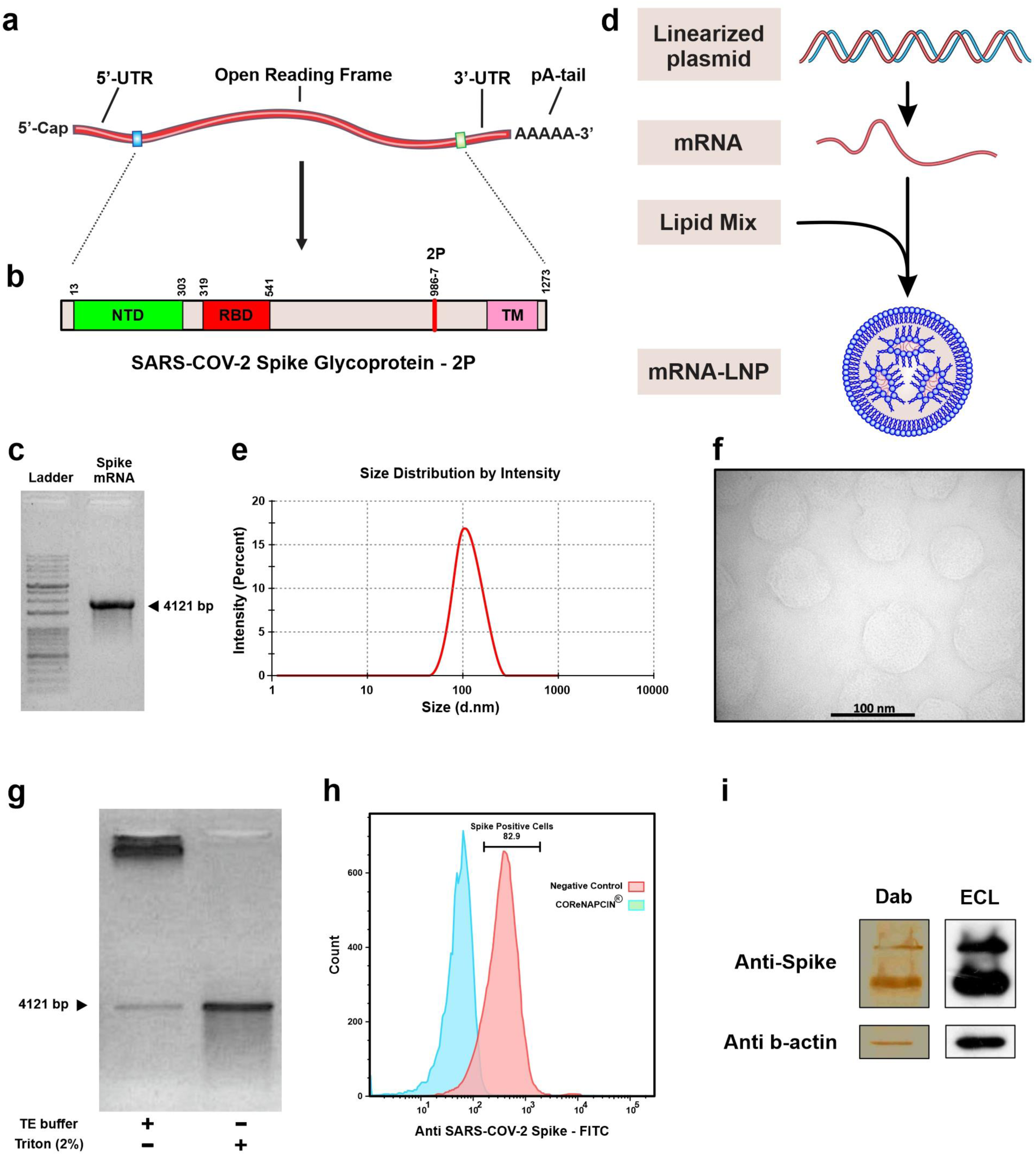
Brief representation of COReNAPCIN^®^ design, production, and evaluation. a) Structure of COReNAPCIN® mRNA. b) Structure of COReNAPCIN^®^ mRNA encoded SARS-CoV-2 Spike glycoprotein-2P. The 2P abbreviation indicates substitution of native amino acids with two proline residues at positions 986 and 987 to produce the pre-fusion stabilized form of the Spike glycoprotein for better antigenic representation at the cell surface. c) Evaluation of the purity and integrity of the produced COReNAPCIN^®^ mRNA with gel electrophoresis. d) Schematic representation of COReNAPCIN^®^ production procedure. The COReNAPCIN^®^ mRNA is made by in-vitro transcription (IVT) from the linearized plasmid DNA template, then is mixed with special lipid components to form the final COReNAPCIN^®^ mRNA-LNP drug product. e) Evaluation of the size distribution of COReNAPCIN^®^ mRNA-LNPs with Dynamic Light Scattering (DLS). f) Evaluation of the COReNAPCIN^®^ particles by Transmission Electron Microscopy (TEM). g) Evaluation of mRNA encapsulation efficiency by Gel Retardation Assay. h) Evaluation of the expression of SARS-CoV-2 Spike glycoprotein-2P at the cell surface by flow Cytometry. i) Evaluation of SARS-CoV-2 Spike glycoprotein-2P expression by Western Blotting.

### 2.2. COReNAPCIN^®^ Immunogenicity in Mice

In order to evaluate the immunogenicity of vaccine candidate, the humoral and cellular immunity were characterized in BALB/c and C57BL/6 mice intramuscularly injected with two doses of 0.05, 0.5 or 3 μg COReNAPCIN^®^ or of PBS, separated by 21 days interval (fig. 2a).

**Figure 2:**
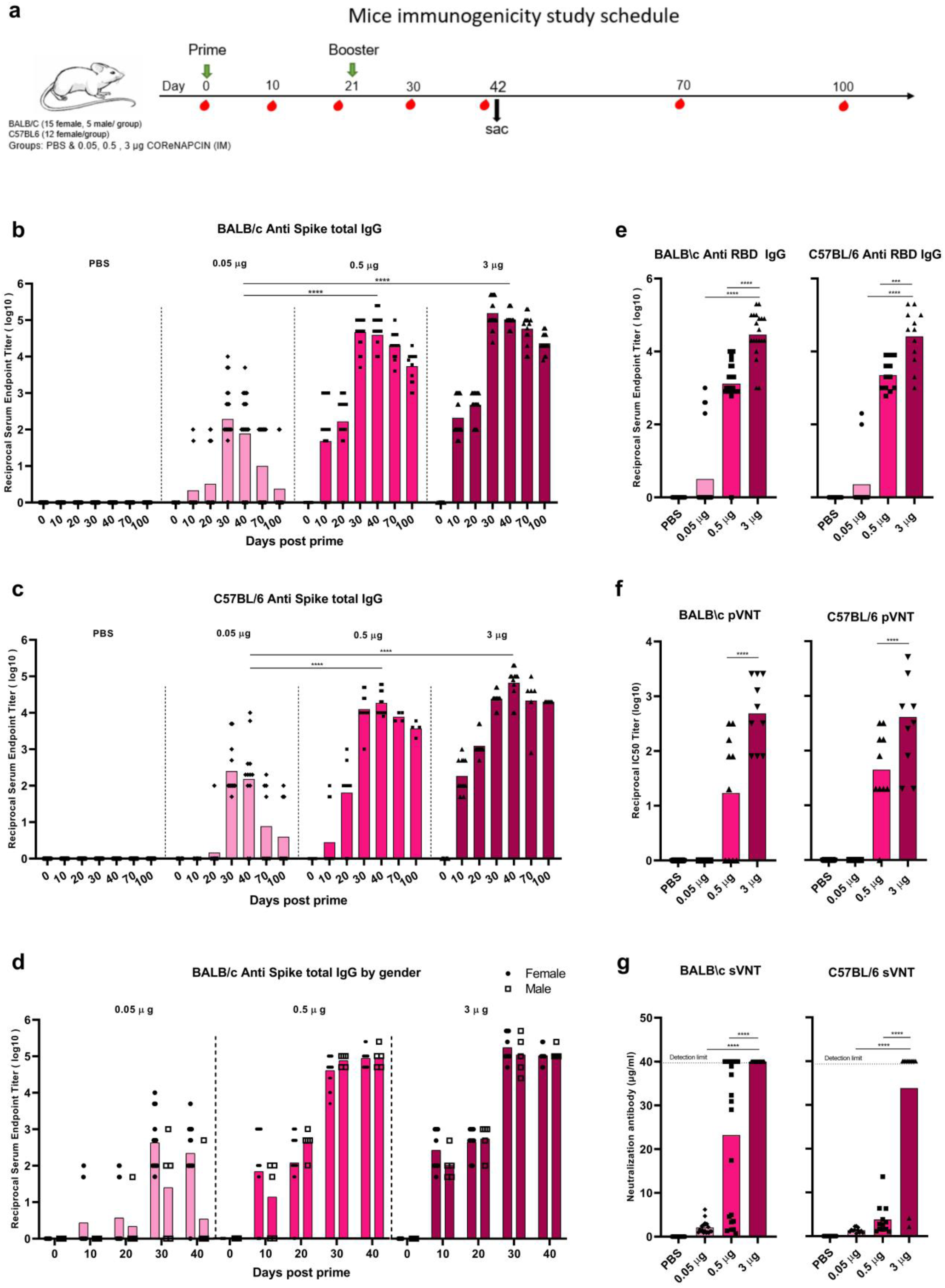
Induction of potent and persistent antibody response with robust neutralizing capacity by COReNAPCIN^®^. a) BALB/c (n= 20 each group) and C57BL/6 mice (n= 12 each group) were i.m. injected with 0.05 (light pink), 0.5 (pink) or 3 μg (purple) COReNAPCIN^®^ at day 0 and day 21. Animals in the control group received PBS injection (gray). b-d) Sera were collected before and at different days post prime and tested for SARS-CoV-2 Spike-specific total IgG by ELISA. In (d), the anti-S IgG specific binding antibody endpoint titer of five different time points in male and female BALB/c is compared. e-g) The sera of day 40 post prime were assessed for anti-RBD IgG binding antibody using ELISA (e), for neutralizing capacity against pseudotype SARS-CoV-2 using pVNT (f) and for ACE2 binding inhibition ability using sVNT (g). A p value less than 0.05 (***P <.001, ****P <.0001) was assumed to be statistically significant.

COReNAPCIN^®^ elicited dose-dependent humoral immune response in both mouse strains (fig. 2). The 0.05, 0.5 and 3 μg doses induced anti-S specific binding antibody which were respectively reached the highest reciprocal Geometric Mean Titer (GMT) of 162.06, 46858 and 154624 in BALB/c and of 250.9901, 18712.47 and 66646.92 in C57BL/6 mice (fig. 2b&c). All BALB/c mice that received more than 0.5 μg vaccine responded to COReNAPCIN^®,^ and the anti-S-specific binding antibody titer showed no statistically significant difference based on gender (fig. 2d). Although the anti-S-specific binding antibody titer decreased over time, it remained at a reasonable level after 100 days post prime (reciprocal GMT of 23028 and 20000, respectively in BALB/c and C57BL/6 mice received 3 μg doses of vaccine candidate). Consistent with anti S specific antibody, the RBD-specific IgG antibody endpoint titer was also elicited by 0.5 and 3 μg of COReNAPCIN^®^ (fig. 2e).

Next, we evaluated whether the COReNAPCIN^®^ elicited antibodies have neutralization potential. For this purpose, serum samples obtained 40 days post prime was assessed by pseudovirus Neutralization Test (pVNT) as well as the ACE2 inhibition kit named as surrogate VNT (sVNT). The serum of mice that received 3 μg COReNAPCIN^®^ showed robust neutralization ability against pseudotyped SARS-CoV-2 in pVNT, with reciprocal GMT of 485 and 411 in BALB/c and C57BL/6 mice, respectively (fig. 2f). Furthermore, as shown in fig. 2g, the neutralization antibody concentration detected by sVNT, reached 40 μg/ml (detection limit of kit) in all mice except two C57BL/6 mice.

There are accumulating lines of evidence which support a role for T cells in immunity against COVID-19^19^. Furthermore, sufficient T cell responses are needed for orchestrating potent antibody responses. Therefore, we analyzed the cellular immune response in mice following the COReNAPCIN^®^ vaccination. Consistent with humoral responses, the induced cellular immunity presented by intracellular cytokine staining and/or ELISpot also exhibited dose-dependency (fig. 3). To explore the S specific T cell responses by intracellular cytokine staining, we stimulated splenocytes from two mice strains with SARS-CoV-2 Spike peptide pool and conducted flow cytometry to quantify the number of Spike specific T cell populations. The frequency of CD44^high^ IFN-γ^+^, CD4^+^ IFN-γ^+^, and CD8^+^ IFN-γ^+^ T cells were significantly increased by 3 μg dose of COReNAPCIN^®^ (fig. 3a-g). T cell responses was further assessed in BALB/c mice using an ELISpot assay. Splenocytes were restimulated with SARS-CoV-2 Spike peptide pool, and IFN-γ-secreting cells were quantified among them. Animals injected with 3 μg COReNAPCIN^®^ showed significantly higher number of Spike specific IFN-γ-secreting cells in response to spike peptide pool (fig. 3h&i).

**Figure 3:**
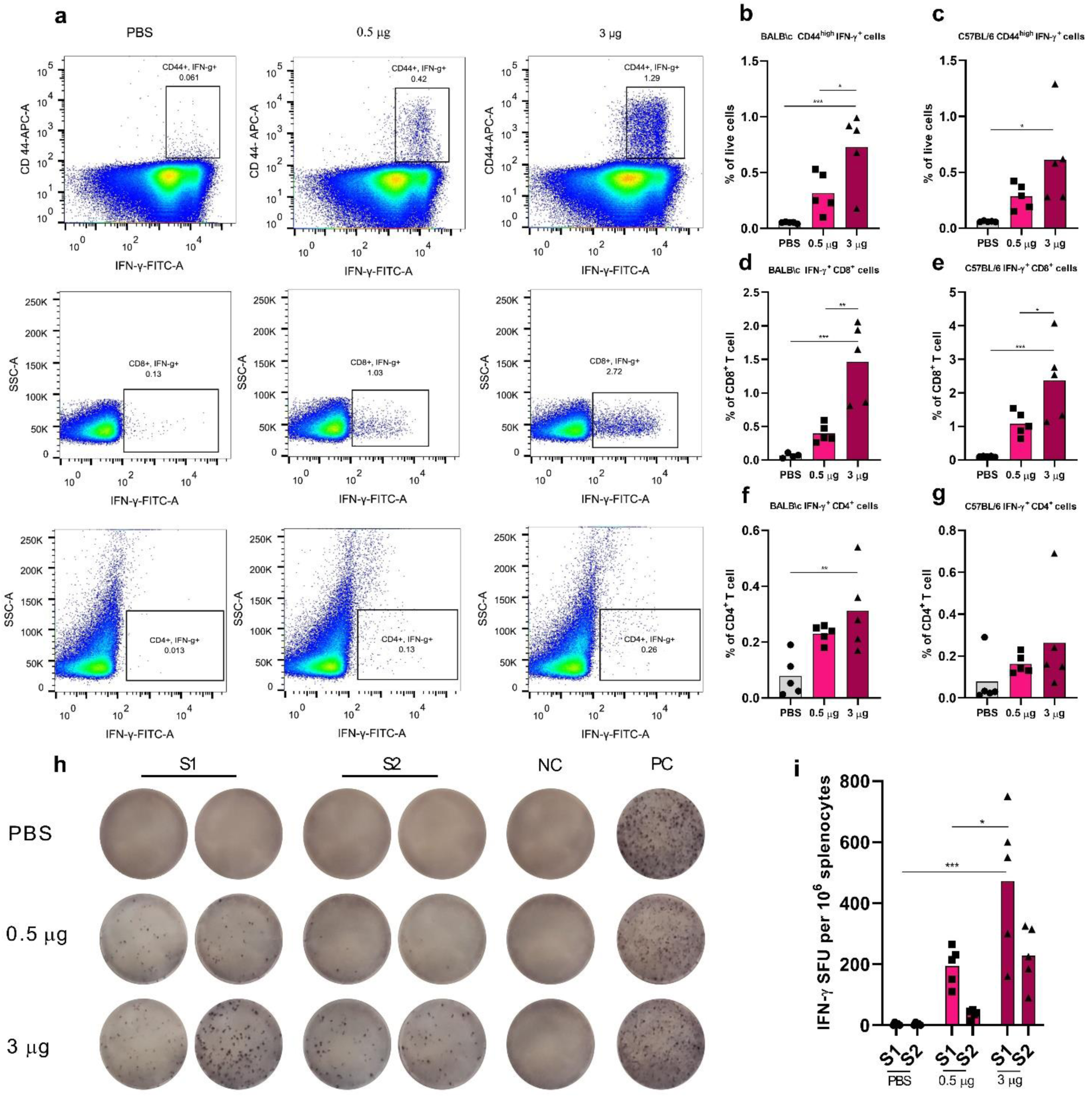
Induction of potent cellular immunity by COReNAPCIN®. Mice were i.m. injected at days 0 and 21 with 0.05 (light pink), 0.5 (pink) or 3 μg (purple) of COReNAPCIN®. Animals in the control group received PBS injection (gray). Splenocytes from five mice each group, were collected at day 21 post boost and subjected for the following assays. a-g) Following stimulation with SARS-CoV-2 Spike peptide pool for 8h, the frequency of Spike specific CD44 ^high^ IFN γ^+^cells (b-c), CD8^+^ IFN γ^+^ cells (d-e) and CD4^+^ IFN γ^+^ cells (f-g) in both mouse strains, were quantified using flow cytometry. Representative dot plots in (a) showing flow cytometry gating summary used for measuring IFN-γ-producing Spike specific T cell populations in C57BL/6 splenocytes. h-i) BALB/c splenocytes were restimulated with two SARS-CoV-2 Spike peptide pool (S1 and S2) for 18 hrs and IFN-γ-secreting T cells were quantified with an ELISpot assay. In (h), the representative images of ELISpot wells, columns from left to right are: responses of splenocytes to S1 SARS-CoV-2 peptide pool (as duplicate), S2 SARS-CoV-2 peptide pool (as duplicate), DMSO (a negative control), and PHA (an unspecific positive control). Each row represents an individual mouse.). A p value less than 0.05 (*P <.05, **P <.01, ***P <.001) was assumed to be statistically significant.

To rule out the possibility of Vaccine-Associated Enhanced Respiratory Disease (VARED), which is a Th2 driven adverse effects following immunization ^20^, we assessed the Th1/Th2 balance in mice. In BALB/c mice, the COReNAPCIN^®^ almost produced the same amount of IgG1 as IgG2a (IgG1 depicts Th2-mediated immunity while IgG2a represents the Th1 phenotype) (fig. 4a&b). Twenty-one days after the boost injection, 5 mice of each group from both strains were sacrificed and their splenocytes were stimulated with a SARS-CoV-2 Spike peptide pool. After 24 hrs stimulation, the secretion of IFN-γ and IL-4 was quantified in the cell culture supernatant. As shown in the fig. 4c&d, there is strong IFN-γ secretion over IL-4, indicating a strong Th1 response over Th2 and thus further confirming that COReNAPCIN^®^ vaccination elicits a Th1-biased immune response.

**Figure 4:**
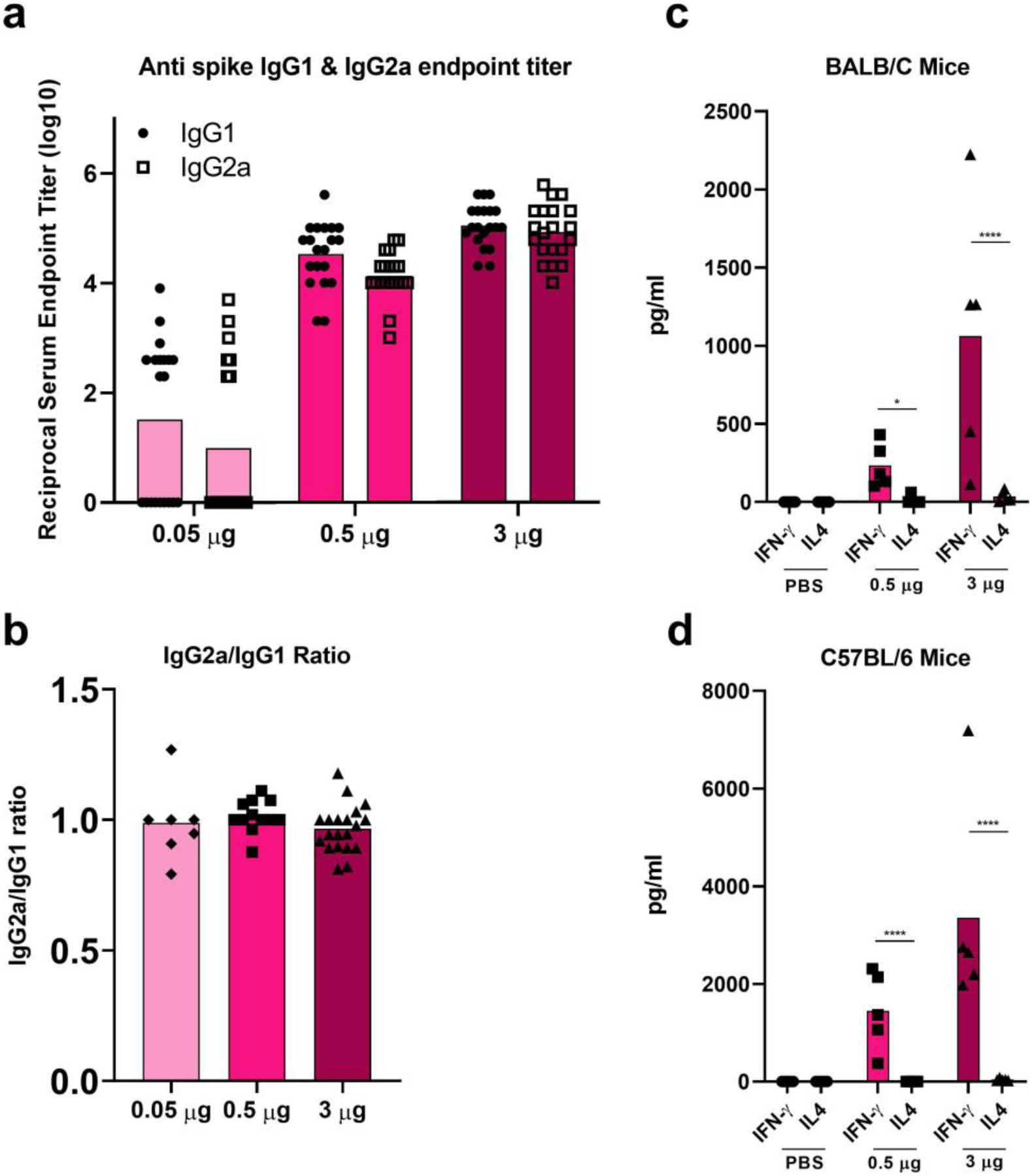
Predominantly Th1-skewed T cell responses by COReNAPCIN^®^. Mice were i.m. injected at days 0 and 21 with 0.05 (light pink), 0.5 (pink) or 3 μg (purple) of COReNAPCIN^®^ or PBS (gray) and the following assays were conducted, 21 days after the booster injection. a-b) Sera from COReNAPCIN^®^ vaccinated BALB/c mice (n= 20 each group) tested for SARS-CoV-2 Spike-specific IgG1 (circle shape symbol, ●) as well as SARS-CoV-2 Spike-specific IgG2a (square shape symbol, □) by ELISA (a) and endpoint titer ratios of IgG2a to IgG1 were calculated (b). c-d) Splenocytes from BALB/c and C57BL/6 mice (n = 5 each group) were exposed to SARS-CoV-2 Spike peptide pool for 24 hrs and the level of IFN-γ and IL-4 cytokines in the cell culture supernatants were determined through ELISA.). A p value less than 0.05 (*P <.05, ****P <.0001) was assumed to be statistically significant.

### 2.3. COReNAPCIN^®^ immunogenicity in non-human primates

Next, we investigated the COReNAPCIN^®^ immunogenicity in rhesus macaques through the same experiments performed in mice. Six rhesus macaques were i.m. injected with two doses of 30 or 50 μg COReNAPCIN^®^ or of PBS with 28 days interval. As shown in the fig. 5b, regardless of the vaccination dose, the anti S specific IgG seroconversion was observed by day 14 after the prime injection and peaked at day 14 post booster dose with mean reciprocal endpoint titer of 1.5×10^6^ and 1.5×10^7^, respectively after the 30 and 50 μg of COReNAPCIN^®^. Although the anti S antibody waned overtime, its reciprocal endpoint titer was remained more than 10^5^ up to day 91 post prime (fig. 5b).

**Figure 5.**
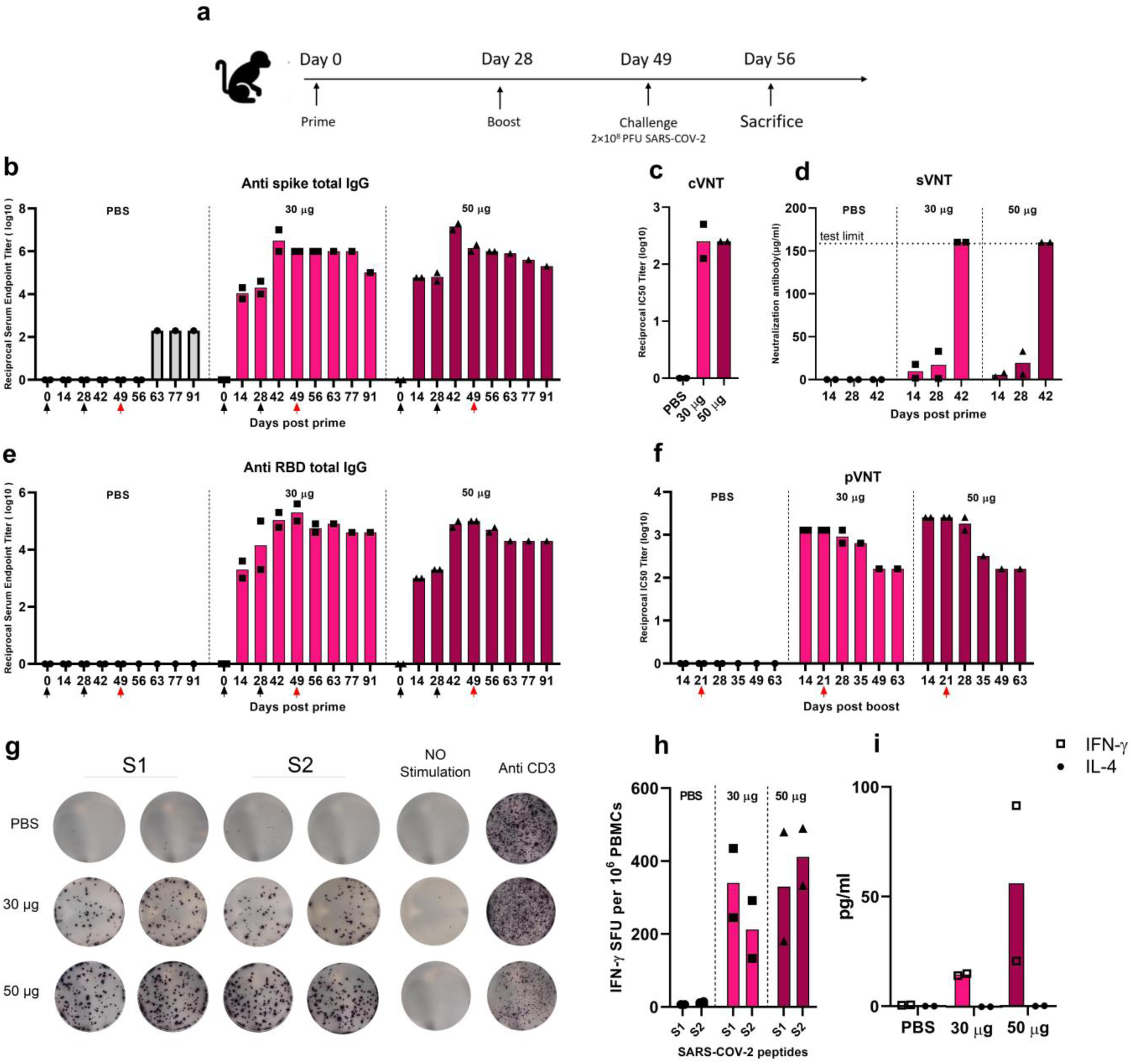
Induction of efficient humoral and cellular immunity in Non-Human Primates by COReNAPCIN®. a) Rhesus macaques (n=2 each group) were immunized with either 30 μg (pink) or 50 μg (purple) COReNAPCIN® at days 0 and 28 and were challenged with 2×10^8^ PFU of SARS-CoV-2, 21 days after the boost injection. Animals in the control group were administrated with PBS (gray). b-f) Serum samples were subjected for evaluating the COReNAPCIN® induced humoral immune responses. The black arrows under the x-axes represent vaccination days while the red one denotes the challenge day. The titer of anti-Spike (b) and anti-RBD (d) specific IgG binding antibody in sera before prime and eight indicated time points post prime were determined by ELISA. The neutralizing capacity were assessed in the sera of day 14 post boost by cVNT (c), and in the sera of indicated different time points by sVNT(d) and pVNT (f). g-i) PBMCs isolated 21 days post-boost injection, were subjected for assessing the COReNAPCIN® induced cellular immune response. Using an ELISpot assay (g-h), after 24 hrs stimulation with SARS-CoV-2 peptide pool (S1 and S2), the frequency of IFN-γ producing cells per 10^6^ splenocytes were quantified (h). In (g), the representative images of ELISpot wells, columns from left to right are: responses of PBMCs to S1 and S2 SARS-CoV-2 peptide pool (as duplicate), DMSO (a negative control), and Anti CD3 (an unspecific positive control). Each row represents an individual rhesus macaque. An ELISA assay (i) was used to assess the IFN-γ (square shape symbol, □) and IL-4 (circle shape symbol, ●) cytokine level in the cell culture supernatants following 8 hrs stimulation with SARS-CoV-2 peptide pool.

We also assessed the anti-RBD-specific IgG binding antibody endpoint titer in rhesus macaques. Same as the anti-Spike antibody, the RBD-specific IgG antibody titer reached its maximum level, 14 days after the booster dose (day 42). As shown in the fig. 5c, we have a 50- to-100-fold increase in anti-RBD antibody endpoint titer 14 days post booster dose compared to 14 days after the prime dose.

Neutralization potential of antibodies elicited against S protein were determined by using a virus-neutralizing test (VNT). Both 30 and 50 μg COReNAPCIN^®^ induced highly potent neutralizing antibodies which have the mean IC50 reciprocal titer of 256, 14 days post-boost (fig. 5c). To further evaluate the potential of SARS-CoV-2 specific antibodies produced in rhesus macaques, we assessed the neutralizing activity of antibodies by pVNT (fig. 5f). The peak for the IC50 reciprocal titer at day 14 post boost reached 2560 for 50 μg and 1280 for 30 μg COReNAPCIN^®^. Consistent with anti S IgG titer, the neutralization titer waned over time, but remained as high as IC50 reciprocal titer of 160 at day 63 post boost. The sVNT test also confirmed the neutralizing activity of antibodies with reaching the detection limit of the test, 14 days post-boost, even with 4-time diluted serum (fig. 5d).

We used ELISpot to assess the cellular immune response in rhesus macaques. At day 21 post-boost, the peripheral blood mononuclear cells (PBMCs) of animals were treated with dimethyl sulfoxide (DMSO) (peptide solvent), SARS-CoV-2 peptide pool, or Anti CD3. The results confirmed that the COReNAPCIN^®^ induces SARS-CoV-2 specific T cells in rhesus macaques at both 30 and 50 μg of the candidate vaccine (fig. 5g&h).

To assess the phenotype of T cell responses elicited by COReNAPCIN^®^ in macaques, we stimulated PBMCs from vaccinated animals with the SARS-CoV-2 peptide pool, and performed enzyme-linked immunosorbent assay (ELISA) assay to quantify the secretion of cytokines in the cell culture supernatants. The results showed strong polarity toward Th1 response over Th2 by the production of predominantly IFN-γ and no IL-4, mainly in rhesus macaques that received 50 μg of candidate vaccine (fig. 5e).

### 2.4. Protection against infection, in rhesus macaques

For evaluating the protective capacity of COReNAPCIN^®^, rhesus macaques, as an infection model for SARS-CoV-2, were subjected to a challenge study in which they were intranasally and intratracheally inoculated with 2×10^8^ PFU of SARS-CoV-2, three weeks after the booster immunization. Nasal and rectal swab were collected at days 2, 4, 7, 14, 21, 28 following infection for determination of RdRp and N genes of SARS-CoV-2. As shown in fig. 6b&c, up to 4 days after the challenge presence of viral transcripts was confirmed in nasal samples from all monkeys. Notably, on day 7 post-challenge, the expression levels of RdRp and N genes of SARS-CoV-2 measured by real-time qRT-PCR, in nasal swabs of COReNAPCIN^®^ immunized macaques showed a prominent decline after day 4 post-challenge. In contrast unvaccinated animals (PBS group) showed significant levels of viral transcripts up until three weeks, with the highest level at day 7 post challenge, representing high viral loads. Expression of SARS-CoV-2 RdRp and N genes in rectal swab specimens of all of the challenged macaques, whether vaccinated or control, showed very high Ct values (fig. S1) indicating no presence of virus.

**Figure 6.**
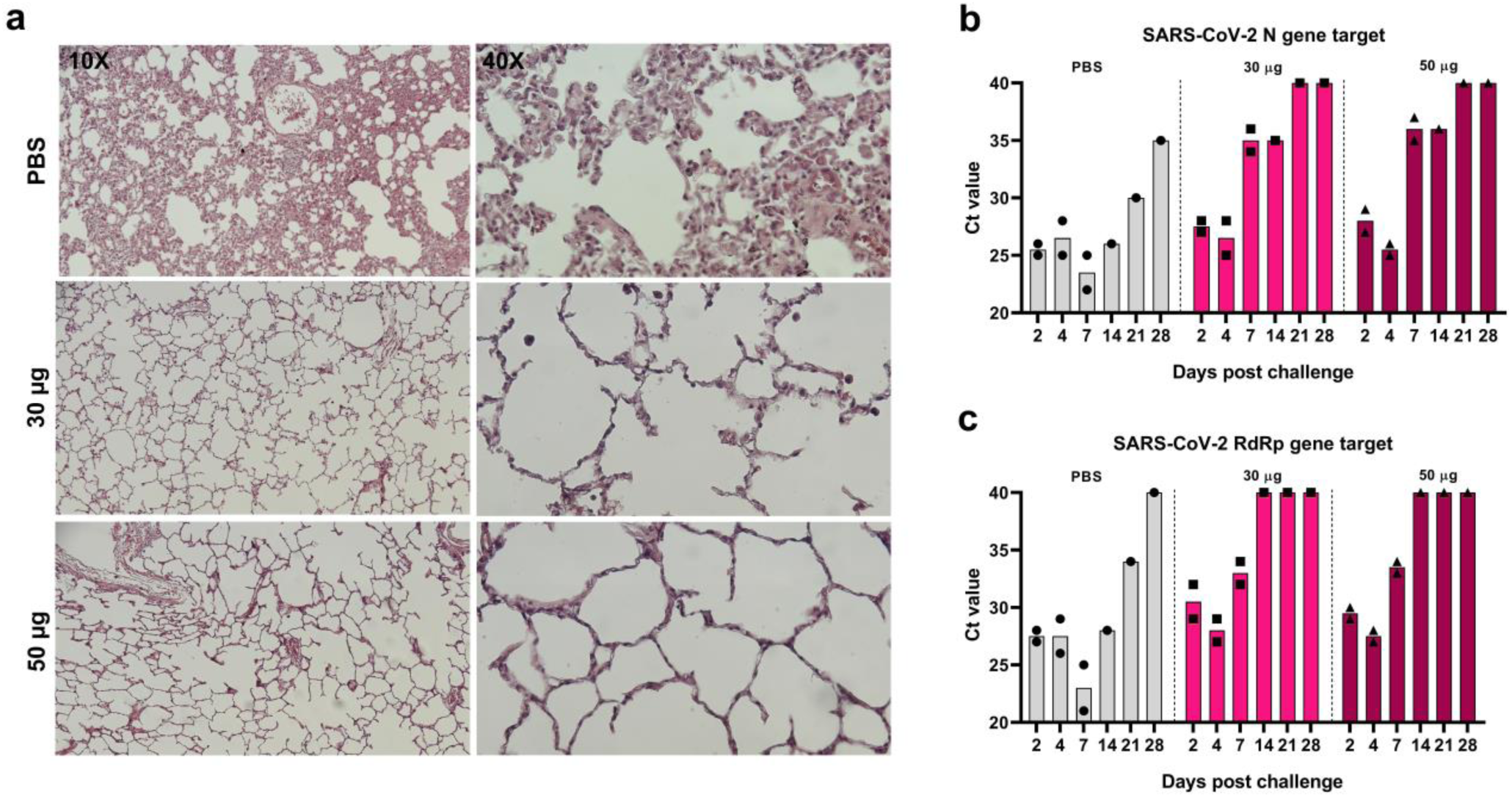
Induction of proper protection against SARS-CoV-2 in rhesus macaques by COReNAPCIN®. Rhesus macaques (n=2 each group) were immunized with PBS or COReNAPCIN^®^ (30 μg or 50 μg) at days 0 and 28 and challenged with 2×10^8^ PFU SARS-CoV-2 on day 21 post boost. a) Seven days post-challenge, lung tissues were harvested from one rhesus macaque per group and were sectioned for hematoxylin and eosin (H&E) staining and histopathology evaluation. The representative images of lung sections from PBS treated and both vaccination groups taken at 10X and 40X magnification are shown. b-c) Nasal swabs were collected at days 2, 4, 7, 14, 21, 28 post challenge, and assessed by RT-PCR for detection of N (b) and RdRp (c) genes of SARS-CoV-2 as indicators of virus presence in the upper respiratory tract.

We further performed histological examination on lung necropsy specimens of three rhesus macaques (one from each group) which were euthanized 7 days post-challenge. There was interstitial pneumonia manifested as thickening of inter-alveolar septa and immune cells infiltration following SARS-CoV-2 infection in unimmunized control animals, while in contrast, both COReNAPCIN^®^ vaccinated animals showed normal pulmonary alveoli structure or very mild histopathological changes with no evidence of considerable inflammation (fig. 6a). This data is in line with the findings of viral genes RT–qPCR, and indicates the protective immunity induction by candidate vaccine. The qPCR and histopathology data support the protective effect of the COReNAPCIN^®^ candidate vaccine in preventing monkeys from being infected with SARS-CoV-2.

## 3. Discussion

After more than two years of COVID-19 experience now billions of people around the world are vaccinated based on different vaccine platforms^1,5^. However, despite the effectiveness of vaccines against ancestral strain, due to the rapid emergence of variant of concerns (VOCs) and also the observed waning immunity, most probably the majority of vaccinated people will require regular booster doses at least in the next few years to overcome this health threatening issue^21–23^. This is true even for people vaccinated with the most effective vaccines; mRNA-1273 & BNT162b, and a similar situation is expected for convalescent people recovered from COVID-19^24–27^. Moreover, due to the high mutation rate and behavior of SARS-CoV-2 circulating between humans and animals, scientists predict COVID-19 will remain a major health challenge probably for the next decade^28,29^. These facts highlight the importance of access to scalable vaccine production platforms for complete eradication of this devastating disease. Therefore, billions of doses will be required annually not only as primary or booster doses for current human population but also for protection of newborns.

As explained earlier, among the different platforms of SARS-CoV-2 vaccines tested so far, mRNA vaccines have some superior unique features. Different studies showed that neutralizing antibody titer of mRNA vaccinated individuals (after 2 doses of vaccination) is somewhere between 8 to 15 times more than convalescent patients, this ratio in inactivated vaccine platforms barely approaches to one^30–32^. Additionally, the reported efficiency of mRNA-1273 and BNT162b, 94.1% and 95% respectively, is among the highest in comparison to other vaccine types^33,34^. Here we have introduced COReNAPCIN^®^; our new mRNA vaccine, revealing outcomes comparable to mRNA-1273 and BNT162b vaccines when administered in mice and rhesus macaques.

In mice, immunization with COReNAPCIN^®^ rapidly elicited high anti-S and anti-RBD binding and neutralizing antibody titers^35,36^. The neutralizing capacity of high titer antibodies induced by COReNAPCIN^®^ were confirmed by pVNT and sVNT. COReNAPCIN^®^ also induced strong T cell responses in mice. Both SARS-CoV-2 specific IFN-γ^+^ CD4^+^ and IFN-γ^+^ CD8^+^ T cells were induced by COReNAPCIN^®^ in mice. Activation of cellular immunity is an important feature of mRNA vaccines especially against new variants, and has also been observed with adenovirus based vaccines^37,38^. However, repeated booster doses of adenoviral vaccines is still under question due to immunity against the vector itself, while mRNA vaccines can be used repeatedly^39,40^. Th1 polarizing vaccines, will reduce the risk of VARED during viral infection^41^. Following COReNAPCIN^®^ injection, a robust Th1-polarized T cell response was observed in BALB/c mice which was confirmed by the balanced IGg2a/IgG1 ratio and IFN-γ production after restimulation with SARS-CoV-2 peptide.

In rhesus macaques, COReNAPCIN^®^ induced strong humoral immunity with the mean neutralizing antibody titer in vaccinated rhesus macaques about 13 folds higher than the titer in convalescent patients (data not shown). COReNAPCIN^®^ also induced a robust cellular immune response with similar immune profile observed in mice. As in mice, Th1 over Th2 response was observed upon vaccination, confirmed by IFN-γ secretion after restimulation with SARS-CoV-2 specific peptide. Following a high dose challenge of macaques with SARS-CoV-2, vaccination with COReNAPCIN^®^ at both 30 and 50μg doses reduced the presence of virus in the upper respiratory tract, and protected rhesus macaques from infection. Challenging the COReNAPCIN^®^ vaccinated macaques with 2×10^8^ PFU SARS-CoV-2; with viral loads about 250 folds higher than that of mRNA-1273 (8×10^5^ PFU) and 95 folds higher than BNT162b (2.1×10^6^ PFU) studies with similar settings showed effective protection in vaccinated vs unvaccinated animals^36,42^. Importantly, the lack of serological response after challenge in vaccinated macaques despite the seroconversion in unvaccinated animals (PBS group), further supports the protection of COReNAPCIN^®^ against SARS-CoV-2 infection.

In summary the preclinical data on immunogenicity of COReNAPCIN^®^, the nucleoside-modified mRNA vaccine against SARS-CoV-2, showed that this new vaccine induces robust humoral and cellular immunity in mice and rhesus macaques and effectively protects rhesus macaques from SARS-CoV-2 infection. Therefore, COReNAPCIN^®^ is expected to show effectiveness comparable to Pfizer & Moderna COVID19 mRNA vaccines in clinical studies. Additionally, COReNAPCIN^®^ variant specific vaccines against major variants of concern are constantly under development and preclinical testing and their outcomes will be presented in future.

## 4. Material and methods

### 4.1. Generation of SARS-CoV-2 mRNA vaccine construct

For mRNA production as the active ingredient of COReNAPCIN^®^ vaccine candidate a genetic construct was designed based on the sequence of Wuhan-Hu-1 SARS-CoV-2 Spike glycoprotein (GenBank: MN908947.3). The codon optimized DNA sequence encoding the full length 1273 amino acid sequence of SARS-CoV-2 Spike glycoprotein was mutated to bear two substitutions (L986P and V987P) for encoding the pre-fusion stabilized form of the protein ^43^. This DNA sequence was cloned into ReNAP-IVT1 plasmid in between special 5’-UTR and 3’-UTR-polyA tail, for transcribing the mRNA under the control of a T7 promoter. Upon sequence verification, the target recombinant plasmid DNA was transformed into E.coli-DH10B bacterial cells and single colonies were selected for preparing COReNAPCIN^®^ vaccine master- and working cell banks (COReNAPCIN^®^-MCB and WCB). After full characterization of master- and working cell banks the WCB was used for the manufacturing step.

### 4.2. COReNAPCIN^®^ mRNA manufacturing

Manufacturing of COReNAPCIN^®^ mRNA starts from one vial COReNAPCIN^®^-WCB. Fed-batch fermentation was performed using a basic defined medium free of animal-derived components. The bacterial pellet was harvested and disrupted by the chemical lysis method with a novel batch operation methodology. Plasmid DNA purification was performed by two-step column chromatography for purification and well separation of plasmid DNA isoforms followed by Ultrafiltration/Diafiltration (UF/DF). The purified circular plasmid DNA was then incubated with restriction enzyme followed by another round of UF/DF to produce linearized plasmid. The ultrapure linearized plasmid was then used as DNA template for producing 5’-capped and polyA tailed COReNAPCIN^®^ mRNA in an in-vitro transcription (IVT) reaction. In addition to linearized plasmid, the IVT reaction was set to contain 5’-cap analog, the mRNA building blocks; ATP, CTP, GTP and N1-Methylpseudouridine-5’-Triphosphate (me1Ψ-UTP), the enzymes; T7 RNA polymerase and inorganic pyrophosphatase, and the RNase inhibitor in a specialized IVT reaction buffer free of animal-derived components. The produced mRNA was then purified quickly with ion exchange (IEX) chromatography followed by oligo-dT affinity chromatography for removing all major impurities such as free 5’-cap and nucleotide analogues, enzymes, plasmid DNA template, as well as degraded, truncated and double-stranded RNAs. The integrity and purity of in-vitro transcribed COReNAPCIN^®^ mRNA was verified before application in the formulation step.

### 4.3. COReNAPCIN^®^ formulation

Following mRNA purification, mRNA was diluted in citrate buffer (pH 4) to reach the desired concentration. Lipid mixture (lipid mix) solution was prepared (12.5 mM concentration) by dissolving a specialized ionizable lipid, cholesterol, DSPC, and DMG-PEG in absolute ethanol with the molar ratio of 50, 38.5, 10, and 1.5, respectively. mRNA and lipid solutions were mixed within a dedicated microfluidic device at the ratio of 3:1 (aqueous: organic phase). Then buffer exchange (acetate buffer, pH 7.4), and concentration processes of formulated mRNA-LNP were performed using tangential flow filtration (TFF). Finally, the formulated mRNA-LNP (COReNAPCIN^®^) was filtrated by 0.22 μm filters and stored at −20°C at the concentration of 0.1 mg/mL.

### 4.4. In vitro characterization of mRNA-LNP

#### 4-4-1 Cell transfection

Primarily, HEK 293T cells were seeded into a 12-well cell culture plate, 2×10^5^ cells per well in HG-DMEM medium (Biowest). After 24 hrs of incubation, cells were transfected with either 1μg mRNA using 2μL Lipofectamine™ (Invitrogen) or 1μg of mRNA-LNP. Untransfected cells were used as negative control. Following 24 hrs of incubation at 37°C, cells were analyzed for SARS-CoV-2 S protein expression by western blot and flow cytometry.

#### 4.4.1. Western blot

Briefly, equal amounts (20μg) of total protein extracts from cultured cells were loaded and run on a 10% SDS-PAGE and then transferred to a nitrocellulose membrane. After 2 hrs blocking in a buffer containing 5% fat-free dried milk and 0.5% Tween-20, the membrane was incubated with anti-SARS-CoV-2 spike polyclonal antibody (Sino biological) or anti-β-actin polyclonal antibody (Abcam) overnight at 4°C. The membrane was then washed three times and incubated for one hour with HRP-conjugated goat anti-rabbit IgG (Sigma) at room temperature. Finally, the result was visualized by DAB staining.

#### 4.4.2. Flow cytometry

In order to analyze transfected cells for cell surface expression of S protein, 2×10^5^ cells of each of transfected and untransfected cells stained with SARS-CoV-2 Spike antibody (SinoBiological) and Anti-human IgG FITC (Sigma-Aldrich®). After staining with primary and secondary antibodies, cells were acquired on BD FACS Lyrics and analyzed by Flowjo V10 (BD Biosciences).

### 4.5. Animal models and immunization

Immunogenicity of COReNAPCIN^®^ vaccine candidate was assessed in mice (BALB/c and C57BL/6) and rhesus macaques. All experiments involving animal, were conducted in accordance with the ethical regulations for the care and use of laboratory animal and approved by Vice-Chancellor in Research Affairs-Tehran University of Medical Sciences with an ethic code of: IR.TUMS.VCR.REC1398.1055. COReNAPCIN^®^ Immunogenicity assessments is summarized in table s1.

#### 4.5.1. Mice immunization

BALB/c and C57BL/6 mice (16-18 g, 4-6 weeks old) were purchased from the Pasteur Institute of Iran. Mice were housed in a standard animal facility with the room temperature of 23 °C, relative humidity of 65% in 12/12-hour light/dark cycles, with free access to water and rodent chow.

Total number of 80 BALB/c mice in four groups (15 females and 5 males per group), and 48 C57BL/6 mice in four groups (12 females in each group), were injected intramuscularly (IM) with either phosphate-buffered saline (PBS) or 0.05, 0.5, or 3 μg mRNA-LNP at days 0 and 21. To evaluate the humoral immune response, blood samples were collected from the retro-orbital sinus of mice under anesthesia. The summary of the mice immunogenicity study design, is presented in fig. 2a. On day 42 post-prime (21 days post-boost), 5 mice from each group (in both BALB/c and C57BL/6 mice) were euthanized, and cellular immune response was assessed by analyzing the splenocytes.

#### 4.5.2. Non-human primate (NHP) immunization

Rhesus macaques were housed individually in cages in a climate-controlled room (temperature of 18–25 °C and humidity 30–70%), with a 12 h light/dark cycles, given chow and fruits in strict accordance to the animal welfare requirements and allowed free access to water. During the challenge study, macaques were housed in the biosafety level 3 (BSL3) facility.

Each group was made of 2 rhesus macaques (1 male and 1 female) injected with either PBS, 30 or 50 μg of mRNA-LNP into the right hind leg (IM) at days 0 and 28. Blood samples were collected and subjected for analyzing humoral immune responses. Cellular immunity was assessed at day 21 post boost injection. The rhesus macaque immunogenicity study design, is illustrated in fig. 5a.

#### 4.5.3. SARS-CoV-2 challenge study of macaques

Rhesus macaques were challenged on day 21 post-boost, under anesthesia, with 2×10^8^ PFU of SARS-CoV-2 in 2 ml PBS, which was divided into two equal volumes for administrating intranasal (1 ml, 0.5 ml per nostril) and intratracheal route (1ml). At days 2, 4, 7, 14, 21, 28 post-challenge, nasal and rectal swabs were collected and analyzed by quantitative Real-time PCR (qRT-PCR) for detection of RdRp and nucleocapsid protein (N) genes of SARS-CoV-2. On day 7 post-challenge, one rhesus from each group was euthanized. After proper fixation of lungs in 10 % buffered formalin, tissue sectioning in 5μm diameter, and hematoxylin and eosin (H&E) staining, lung sections were analyzed by a veterinary pathologist.

### 4.6. Quantification of SARS-CoV-2 RNA by qPCR

To detect and measure the presence of SARS-CoV-2 viral RNA in nasal turbinates and rectal swab samples of macaques, two SARS-CoV-2 genes, N and RdRp genes, were analyzed by qPCR test, using SENMURV multi-star SARS-CoV-2 RT-PCR kit (STRC). Nasal and rectal swab specimens were subjected to total RNA extraction using the RNeasy Mini Kit (QIAGEN). The qRT-PCR reaction, was run with the following thermal profile: 20 min in 50°C, 10 min in 95°C, and followed by 40 cycles of 95°C and 55°C for 10 and 40 s, respectively.

### 4.7. Enzyme-linked immunosorbent assay (ELISA)

SPL Maxibinding ELISA plates (SPL) were coated with 33 ng of SARS-CoV-2 S protein (Acro-Biosystems) or SARS-CoV-2 receptor binding protein (RBD) (Sino Biological) in 100 μl (final concentration of 330 ng/ml) coating buffer (Biolegend) with overnight incubation at 4°C. After routine wash and block with 2% Bovine Serum Albumin, (BSA) (Sigma-Aldrich), 100 μl of serial diluted mice or macaques serum were added to wells and incubated at room temperature for one hour. After washing, 50μl of either Anti-Mouse IgG (γ-chain specific), Anti-mouse IgG1, Anti-mouse IgG2a (Sigma-Aldrich, USA) or Anti-monkey IgG in 1:5000 dilutions were added to each well and plates were incubated at room temperature for one hour.

After 4 rounds of wash, 100 μl of Tetramethylbenzidine (TMB) (RaziBIOTech) were added to each well, followed by 10 min of incubation. Finally, 1N HCl was used as a stop reagent and the absorbance was measured by BioTek^®^ 800 ™ TS at 450 nm and 630 nm. To calculate endpoint titers, the baseline serum samples of 30 mice, were used and calculated as described by Frey *et*.*a*l^44^.

### 4.8. Pseudovirus neutralization assay (pVNT)

Spike pseudotyped lentivirus, produced by the co-transfection of plex307-egfp, pCMV3-spike (Wuhan-D614G), and pspax2 with Lipofectamine 3000 (Thermo). Spike pseudotyped lentivirus encodes the EGFP protein that serves as a reporter gene. pVNT test performed as previously described by Ferrara *et*.*al* ^45^. Briefly, sera was diluted 2-fold serially from 1/40 in a 96-well cell culture plate. After one hour incubation with Spike pseudotyped lentivirus in 37°C, 14×10^3^ of HEK-293T-hACE2 cells were added to each well. Fluorescent microscopy was used to count the number of EGFP-positive cells of each well, after 48 to 60 hrs. IC50 was calculated using the percentage of GFP positive cell versus log (dilution factor) in GraphPad Prism V8.

### 4.9. Conventional virus neutralization test (cVNT)

The cVNT assay, was conducted on serum samples of rhesus macaques, collected on day 14 post boost. Briefly, 50 μL of 100 median tissue culture infecting dose per mL (TCID_50_/mL) of SARS-CoV-2, were mixed with 50 μL of two-fold serial dilutions of heat-inactivated serum (at 56°C for 30 min). After 60 min incubation at 37°C, the virus-serum mixture was added to 96 well-plate previously seeded with 1.5 × 10^4^ Vero cells per well (performed in triplicate). One hour post-incubation, the supernatant was removed and cells were washed twice with Dulbecco’s Modified Eagle Medium (DMEM). Then, cells were incubated for 72 hrs at 37°C in a 5% CO2 in DMEM with 10% heat-inactivated Fetal bovine serum (FBS). Half-maximal inhibitory concentration (IC50) titer, was measured by microscopy and reported as the highest dilution inhibiting 50% cytopathic effect (CPE) formation.

### 4.10. Surrogate virus neutralization test (sVNT)

We used sVNT (ACE2 inhibition test) ELISA (Pishtaz Teb) (with a detection limit of 40 μg/ml), for measuring the SARS-COV-2 neutralizing ability of serum samples of mice and rhesus macaques collected 40 days post-prime. The assay was performed according to the manufacturer’s instructions; in summary, 50μl of serum samples, negative and positive controls were added to the wells. Then immediately, 50μl RBD conjugated HRP, was added and incubated for 30 min at 37°C. Following the routine wash, 100 μl TMB was added to each well and plates were placed at room temperature for 15 min. Finally, 100 μl 1N HCL as stop solution was added to each well and the absorbance was measured by BioTek® 800 ™ TS at 450 nm and 630 nm (as Reference filter).

### 4.11. Intracellular cytokine staining

Mice fresh splenocytes were separated by mechanical homogenization then filtered through a 70 μm cell strainer (Biologics). Red Blood Cells (RBCs) were removed by using RBC lysis buffer, then following wash with PBS, 10^6^ splenocytes were ex vivo stimulated with SARS-CoV-2 peptide pool (JPT PM-WCPV-S-1) at concentration of 2 μg/ml of each peptide in RPMI (Biowest) supplemented with 10% heat inactivated FBS (Gibco). PMA/Ionomycin (BD Pharmingen™) and dimethyl sulfoxide (DMSO) (Sigma-Aldrich) were used as a positive and negative control, respectively. Brefeldin (Biolegend) as Golgi stop was added to each well one-hour post-incubation. Following eight hrs of incubation, the harvested cells, washed with PBS, and stained with Zombie Violet (Biolegend) at 1/200 dilution and incubated at room temperature for 30 min. After washing cells with staining buffer (PBS supplemented with 2% FBS and 0.05% NaN3), anti CD3-Percp, anti CD8-PE, anti CD4-PE, anti CD44-APC (Biolegend) were used for cell surface staining followed by 20 min incubation at room temperature. For intracellular staining, cells were fixed and permeabilized with Cytofast fix/perm (Biolegend) according to the manufacturer instruction, then the anti IFN-γ-FITC (Biolegend) was added to the tubes. After 20 minutes incubation and washing with permeabilization and staining buffer (Biolegend), cells were acquired on BD FACS Lyrics and analyzed by Flowjo V10 (BD Biosciences).

### 4.12. Enzyme-linked immunosorbent spot assay (ELISpot)

Cryopreserved mice splenocytes were thawed in pre warmed RPMI (Biowest) and rested for 24 h. IFN-γ ELISpot precoated plates (Abcam) were used according to the manufacturer’s instruction. Briefly, the total number of 2×10^5^ harvested splenocytes, were stimulated with 2 μg/ml of SARS-CoV-2 peptide pool (JPT product PM-WCPV-S-1, Germany), for 18 hrs at 37° C. DMSO (Sigma-Aldrich, USA) and phytohemagglutinin (PHA) (Gibco) were used as negative and positive control, respectively. After several wash steps, 100 μl streptavidin-AP (Abcam) was added to each well, followed by incubation at room temperature for 1 hour. Finally, 100 μl BCIP/NBT was added to the each well and after 15 min incubation at room temperature, plate was rinsed in tap water, dried and spots were counted by image processing software.

For quantitative measuring of macaque specific T cell, Monkey IFN-γ ELISpot plates (Mabtech) were used. After thawing and overnight incubation, total number of 3×10^5^ NHP PBMC, were added to each well and stimulated with 2 μg/ml of SARS-CoV-2 peptide pools (JPT, PM-WCPV-S-1). DMSO (peptide solvent) and anti CD3 (Mabtech) used for negative and positive control, respectively. The plate then incubated at 37°C for 18 hrs followed by wash according to the manufacturer instruction. After incubation with detection antibody (7-B6-ALP) and subsequent washing, 100 μl substrate solution (BCIP/NBT-plus) added to each well. When spots were formed, plate rinsed in tap water, dried and spots were counted by image processing software.

### 4.13. Cytokine analysis

Splenocytes and peripheral blood mononuclear cells (PBMCs), respectively, from immunized mice and rhesus macaques were collected 21 days post boost. Five-million mice splenocytes or rhesus macaque PBMCs were stimulated for 24 hrs with SARS-CoV-2 peptide pool (JPT, PM-WCPV-S-1) at final concentration of 2 μg/ml per peptide. Negative control wells received same volume of DMSO with no peptide and PMA/Ionomycin added to positive control wells. Concentrations of IFN-γ and IL-4 in supernatants were analyzed by DuoSet ELISA kit (Biolegend) according to the manufacturer’s instructions.

### 4.14. Statistical analysis

Graph Pad V8 Prism was used to illustrates the figures and analyze the results statistically. Dotted lines indicate assay limits of detection. For comparing values between two groups, t-test was conducted and Group comparisons were made by one-way ANOVA followed by Dunn’s test. A p value less than 0.05 (*P <.05, **P <.01, ***P <.001, ****P <.001) was assumed to be statistically significant.

## 5. Acknowledgment

We acknowledge Vice-presidency for science and technology (VPST), Biotechnology Development Council (BDC), Council for Development of Stem Cell Sciences (IRSCC) and Technologies and Tehran University of Medical Sciences (TUMS) for their extraordinary support. The authors also gratefully acknowledge the financial support of Iran Biotech Fund (BIF), and Iran National Innovation Fund (INIF). Most of this work was carried out within the Stem Cell and Regenerative Medicine Center of Excellence at TUMS and TUMS Preclinical Core Facility (TPCF). We thank their consistent, warm and kind cooperation for providing excellent infrastructure for conducting experiments. Finally, we are immensely grateful to work with PersisGen Par & AryoGen Pharmed companies for their contribution in GMP-grade manufacturing of COReNAPCIN^®^.

## 7. Author contributions

The authors V.K., S.D., V.S., S.M.H. designed the study. S.D., V.K and K.N designed the vaccine genetic constructs. T.M made the genetic constructs. D.S.K. made MCB and WCBs. K.A., and M.S.M. produced and purified plasmid and mRNA at small scales. M.P., and M.D. developed the high-throughput plasmid and mRNA production and purification platform. M.P., M.D., D.S.K., M.H.G., H.R., H.E.B., and H.B. conducted high-throughput plasmid and mRNA production and purification. M.A. and M.P. designed and made the high-throughput plasmid production devices. L.R., and S.M.S.S. with the help of M.H.M. developed the ionizable lipid for LNP formulation. R.R., S.H.K., M.H.M., S.D, and V.K. designed the LNP formulation. R.R., E.A.D., S.H.K., F.M.P., S.S.M. with the help of M.S.N. and M.P. conducted the LNP formulation. S.A.M.S., R.R. and V.K. designed the microfluidic system for LNP formulation. S.A.M.S. with the help of S.A. and H.R.N. developed the microfluidic device and microchips for LNP formulation. V.K, S.D, F.N., M.E., L.R, M.S.M. designed the quality control tests. M.E., F.N., F.H., F.M.P., L.R., N.M. with the help of K.N., M.S.M. and P.P. performed the quality control tests. M.E., K.N. designed the pseudotype virus neutralization test (pVNT) and with the help of V.T. conducted the pVNTs. R.M., and M.H.M.A established the production line for IVT enzymes and the RNase inhibitor and A.M., S.S.M., A.H.R., M.R. with the help of F.B., and M.A. produced the proteins. S.M.H., S.D. with the help of V.S. designed the preclinical studies. R.A., M.H., S.P., M.A. conducted the preclinical tests. R.A., M.H., S.P., M.A, M.P., S.H.K., M.E. and V.K. wrote the manuscript. R.A., M.H., S.P., M.A., and V.K. produced the figures. S.D., S.M.H., V.S., S.H.K., M.S.M and V.K. edited the manuscript.

