## Supplementary material for "mRNA-based vaccine candidate COReNAPCIN^®^ induces robust humoral and cellular immunity in mice and non-human primates": Fig. s1

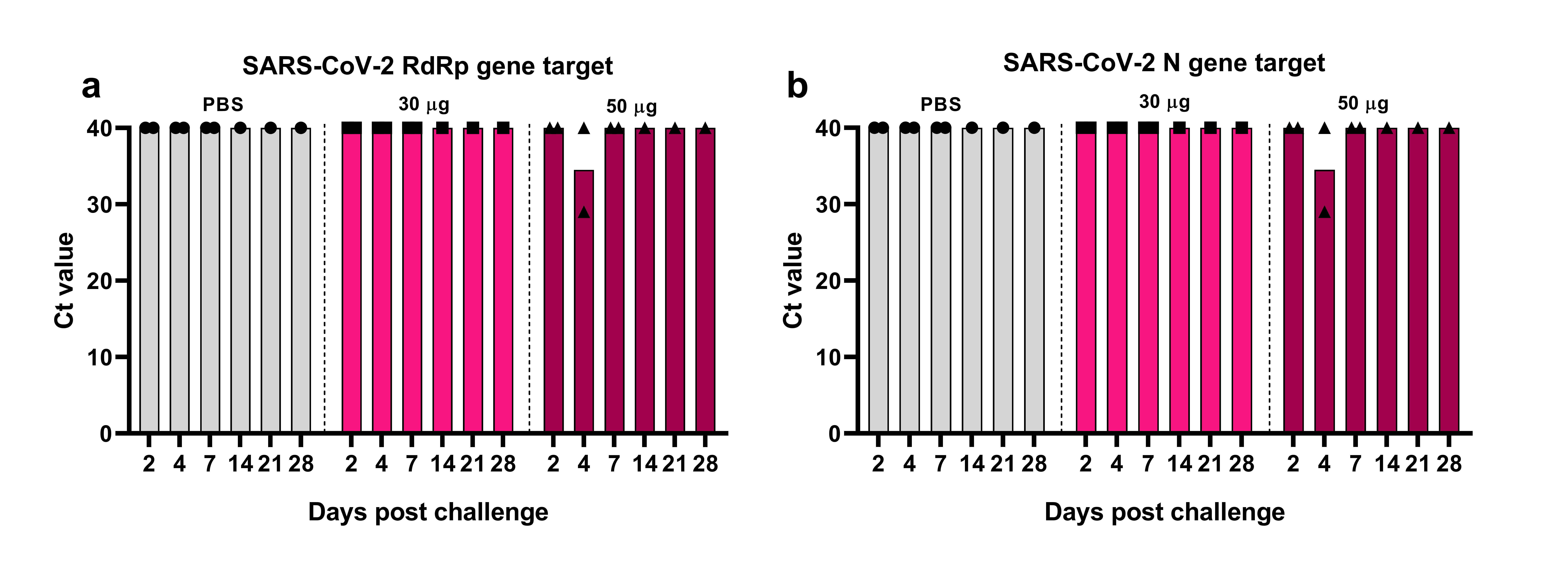

**Figure S1: Assessment of SARS-CoV-2 presence in rectal swabs.** *Rectal swabs were collected at days 2, 4, 7, 14, 21, 28 post challenge, and assessed by RT-PCR for detection of RdRp (a) and N (b) genes of SARS-CoV-2.*
