## Supplementary material for "mRNA-based vaccine candidate COReNAPCIN^®^ induces robust humoral and cellular immunity in mice and non-human primates": Table s1

**Table s1: COReNAPCIN^®^ Immunogenicity assessments summary.**

| **Test** | | **BALB/C Mice** | **C57BL6 Mice** | **Rhesus Macaque** |
| --- | --- | --- | --- | --- |
| **Humoral immunity** | | | | |
| **Anti-Spike specific IgG binding antibody titer** | | ✓ | ✓ | ✓ |
| **Anti-RBD IgG binding antibody titer** | | ✓ | ✓ | ✓ |
| **cVNT** | | - | - | ✓ |
| **pVNT** | | ✓ | ✓ | ✓ |
| **sVNT(ACE2 inhibition)** | | ✓ | ✓ | ✓ |
| **Th1/Th2 balance** | | | | |
| **IgG1 /IgG2 ratio** | | ✓ | - | - |
| **SARS-CoV-2 specific cytokines secretion by**  **Splenocytes/PBMCs** | IFN-γ | ✓ | ✓ | ✓ |
|  | IL-4 | ✓ | ✓ | ✓ |
| **Cellular immunity** | | | | |
| **SARS-CoV-2 Spike specific T cell populations (flow cytometry)** | | ✓ | ✓ | - |
| **ELISPOT (SARS-CoV-2 Spike-specific IFN-γ-secreting Splenocytes/PBMCs)** | | - | - | ✓ |
| **Challenge study** | | | | |
| **2×10^8^ PFU of SARS-CoV-2 Challenge** | | - | - | ✓ |
| **Virus detection**  **(PCR for detection of nasal and rectal RdRp and N genes )** | | - | - | ✓ |
| **Frequency of PMNs in BAL**  **(cell counting and differential analysis)** | | - | - | ✓ |
| **Histopathology after challenge** | | - | - | ✓ |
